# Non-Consummatory Behavior Signals Predict Aversion-Resistant Alcohol Drinking in Head-Fixed Mice

**DOI:** 10.1101/2023.06.20.545767

**Authors:** Nicholas M. Timme, Cherish E. Ardinger, Seth D. C. Weir, Rachel Zelaya-Escobar, Rachel Kruger, Christopher C. Lapish

## Abstract

A key facet of alcohol use disorder is continuing to drink alcohol despite negative consequences (so called “aversion-resistant drinking”). In this study, we sought to assess the degree to which head-fixed mice exhibit aversion-resistant drinking and to leverage behavioral analysis techniques available in head-fixture to relate non-consummatory behaviors to aversion-resistant drinking. We assessed aversion-resistant drinking in head-fixed female and male C57BL/6J mice. We adulterated 20% (v/v) alcohol with varying concentrations of the bitter tastant quinine to measure the degree to which mice would continue to drink despite this aversive stimulus. We recorded high-resolution video of the mice during head-fixed drinking, tracked body parts with machine vision tools, and analyzed body movements in relation to consumption. Female and male head-fixed mice exhibited heterogenous levels of aversion-resistant drinking. Additionally, non-consummatory behaviors, such as paw movement and snout movement, were related to the intensity of aversion-resistant drinking. These studies demonstrate that head-fixed mice exhibit aversion-resistant drinking and that non-consummatory behaviors can be used to assess perceived aversiveness in this paradigm. Furthermore, these studies lay the groundwork for future experiments that will utilize advanced electrophysiological techniques to record from large populations of neurons during aversion-resistant drinking to understand the neurocomputational processes that drive this clinically relevant behavior.

## 1 Introduction

The majority of the criteria used to assess the severity of an alcohol use disorder (AUD) involve continuing to drink despite suffering negative consequences [1]. Thus, continuing to drink alcohol despite negative consequences (so called “aversion-resistant drinking” (ARD)) is a key feature of AUD. Furthermore, one possible avenue to develop new treatments for AUD is to identify and repair the maladaptive neural processes underlying ARD. In this study, we describe a series of experiments that characterized ARD in head-fixed mice and leveraged behavior analysis techniques available in head-fixture to relate non-consummatory behaviors to ARD.

A standard method for modeling aversion-resistant drinking in rodents is to assess the degree to which an animal is willing to consume alcohol that is adulterated with the bitter tastant quinine [2-6], with the view that animals that are more willing to consume quinine adulterated alcohol are more aversion-resistant. This method of assessing ARD has been used in numerous animal models, including mice [6-17], Wistar rats [18-24], alcohol preferring P rats [18-20, 25, 26], and Sprague-Dawley rats [27].

Prefrontal cortex (PFC) is known to play a key role in addiction [28-30] in general, but it has been directly implicated in ARD in particular. Bidirectional modulation of projections from medial PFC to dorsal periaqueductal gray have been shown to bidirectionally affect ARD in mice [10] and differences in neural activity as measured by FOS activity has been observed in male and female mice during ARD in ventromedial PFC [7]. Medial PFC neural populations in mice encode information about and can regulate operant responding during footshock punished drinking [31]. Furthermore, numerous other brain regions that interact with PFC, such as the ventral tegmental area [7], the dorsal striatum [11], the nucleus accumbens [15], the amygdala [7, 27, 32, 33], and the insular cortex [7, 16, 34] have also been implicated in ARD using various rodent models. Taken together, these results suggest that altered interactions among the PFC and the numerous brain regions it shares connections with may be crucial for the expression of ARD. Testing this hypothesis requires the use of techniques that are capable of recording neural behavior across multiple brain regions in order to understand the neurocomputational phenomena underlying ARD.

Head-fixed mouse preparations offer the potential to meet this need because these methods allow researchers to exploit powerful neural recording techniques, such as calcium imaging [35, 36] and high-yield, multi-brain region electrophysiology [37, 38]. Head-fixed methods have been widely utilized in various types of non-addiction related studies [39-41]. For instance, these techniques make it possible to record from large portions of the cortical surface using wide-field calcium imaging [42], to simultaneously record from and manipulate brain region level populations with two-photon calcium imaging and optogenetics [43], or record from thousands of neurons simultaneously across multiple brain regions using multiple high-density electrophysiology probes [44]. Furthermore, due to the stable positioning of the animal, these methods facilitate motion tracking of individual body parts [45] and advanced behavioral analyses [46-55], as well as the precise delivery of visual [56], auditory [57], and olfactory stimuli [58, 59]. This improved precision reduces noise between behavioral data, stimuli, and neural activity in comparison to freely moving mice. Despite the advantages of these head-fixed techniques, as well as extensive research on alcohol consumption and ARD using quinine in freely moving mice, very little data exist regarding alcohol consumption and ARD in head-fixed mice. To date, it has been shown that head-fixed mice will voluntarily consume alcohol [46], but no studies have documented ARD using quinine in head-fixed mice. Furthermore, no studies have related head-fixed alcohol drinking and ARD to home-cage drinking patterns and ARD. A growing body of literature has highlighted the importance of alcohol drinking pattern – especially alcohol frontloading wherein animals drink large quantities of alcohol quickly – as a behavioral readout of the animal’s motivation to consume alcohol [60]. Therefore, relating home-cage drinking patterns to head-fixed drinking has the potential to improve our understanding of drinking motivation in head-fixation.

In this study, we sought to address this void in the literature by examining ARD in head-fixed mice. We hypothesized that a subset of head-fixed mice would exhibit ARD, mice that tended to frontload in home-cage would exhibit high degrees of ARD in head-fixture, and that non-consummatory behaviors would be related to ARD. These studies will lay the groundwork for future neural recordings during ARD in head-fixed mice.

## 2 Materials and Methods

### 2.1 Subjects

Female (N = 13) and male (N = 13) C57BL/6J mice (Jackson Labs) were used in this study. Two females and one male were lost due to failed head bar implants and were excluded from the study. Experiments were conducted in two cohorts (cohort 1: 6 females and 5 males, cohort 2: 5 females and 7 males). All animal procedures were approved by the Indiana University – Purdue University Indianapolis School of Science Institutional Animal Care and Use Committee.

### 2.2 Surgery

At six weeks of age (Figure 1 A), mice underwent head-bar implantation surgery. We utilized a modified version of the International Brain Laboratory surgery protocol [61]. Mice were anesthetized using isoflurane and placed in a stereotaxic frame (Kopf Instruments). Mice were administered cefazolin intraperitoneally (i.p.) at a dose of 30 mg/kg. The animal’s scalp was sterilized using betadine and alcohol wipes. The scalp was anesthetized using an injection of bupivacaine into the scalp at a dose of 2.5 mg/kg prior to removal. A layer of Optibond (Kerr) was applied to the cleared skull and cured with UV light. The skull was leveled and the headbar was positioned along the midline of the skull, approximately 0.1 mm above the skull with the posterior side of the bar positioned at -2.95 mm A/P from bregma. The bar was cemented to the skull using C&B Metabond (Parkell). The cement was allowed to dry for approximately 10 minutes, during which Ketofen was administered i.p. at a dose of 5 mg/kg. The animal was then removed from the stereotax and allowed to recover in a warmed chamber until it regained righting reflex. The animal’s health and weight were monitored for 7 days post-surgery for signs of infections or complications.

**Figure 1:**
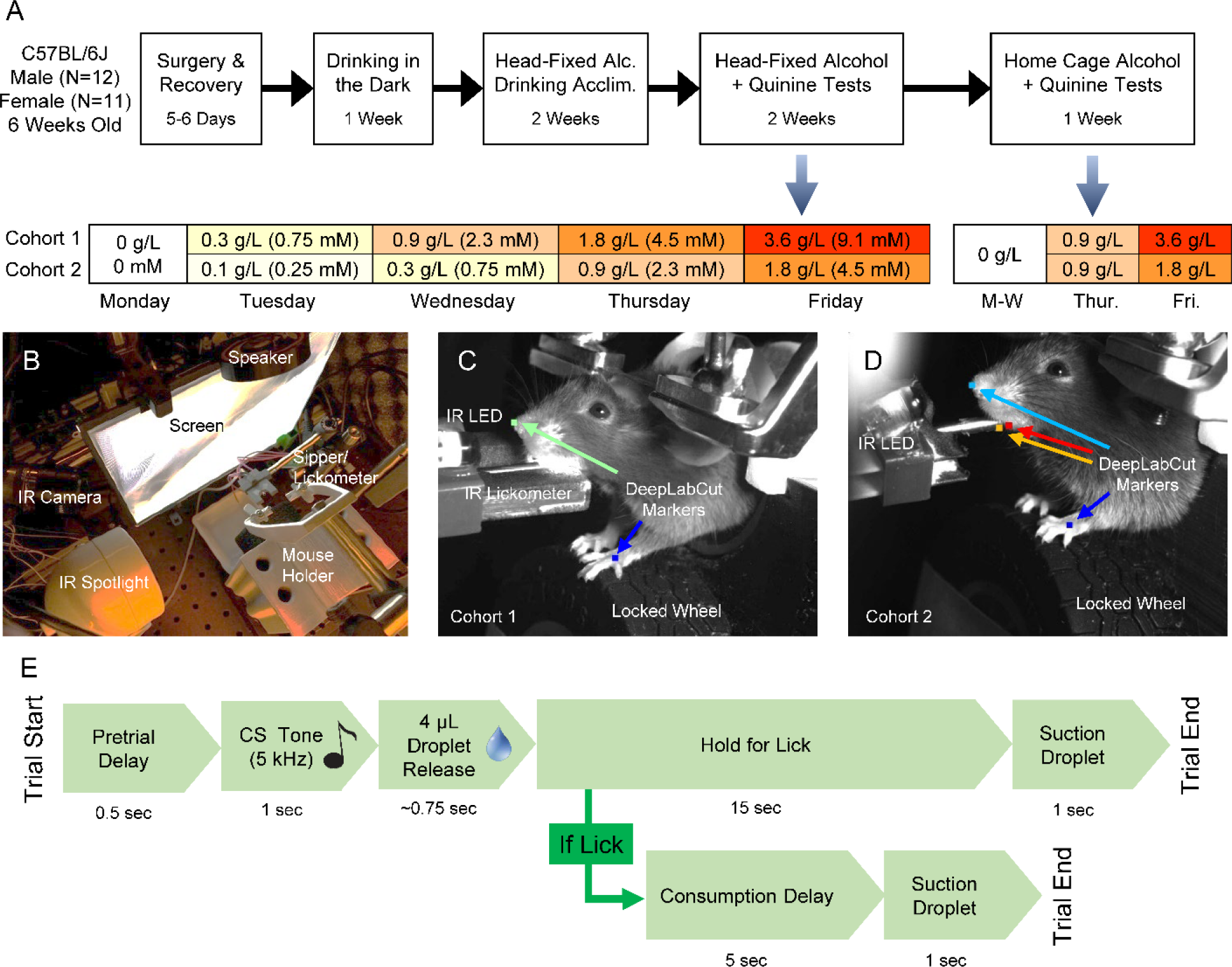
Experiment schedule, trial sequence, and head-fixation apparatus. **(A)** Experiment schedule. Two cohorts were used (1: 5M, 6F; 2: 7M, 5F). Mice underwent head-bar implantation surgery followed by one week of DID. Then, mice were acclimated to head-fixed alcohol drinking over 2 weeks. Next, mice underwent two weeks of quinine adulterated drinking with increasing concentrations of quinine used each week (i.e., two ramps). Finally, mice underwent a home-cage quinine adulterated alcohol drinking test. **(B)** View of head-fixed rig from above. **(C)** Image of a head-fixed mouse from cohort 1. An IR lickometer was used to record licks and two body points were tracked to assess non-consummatory behavior. **(D)** Image of a head-fixed mouse from cohort 2. The IR lickometer was replaced with a real-time machine vision image classification system to assess licking. Four body parts were tracked to assess non-consummatory behavior and post-hoc analysis of licking. **(E)** The head-fixed cued access task consisted of a brief pretrial delay, followed by a 1 sec conditioned stimulus tone (5 kHz), followed by the release of a 4 uL droplet onto the end of the sipper. The animal was given 15 seconds to lick the droplet. If no lick was recorded, the droplet was suctioned off the sipper. If a lick was recorded, the animal would receive 5 seconds to finish consuming the droplet before it would be suctioned off. Each test took approximately 30 minutes.

### 2.3 Drinking in the Dark (DID) and Frontloading

Approximately 1 week after surgery, mice underwent 1 week of home-cage drinking using the drinking in the dark (DID) paradigm (Figure 1 A) [62]. Mice received access to 20% ethanol (v/v) (Decon Laboratories, Inc.) and no water for 2 hours a day, Monday through Friday. Mice were single-housed in standard shoebox cages in a room with a 12-hour reverse light/dark cycle. Ethanol access was timed to 3 hours into the mice’s dark cycle (noon to 2 PM). Animals had ad libitum access to regular lab chow (LabDiet 5001) [63] at all times when not head-fixed. Ethanol during DID was made available to mice in volumetric sipper tubes (Columbus Instruments, Inc.). These sippers measured the volume of consumed ethanol each minute, which allowed for the analysis of drinking patterns to assess if there was evidence for the presence of frontloading behavior. This approach has been used previously [63-66].

We used recently introduced techniques to identify alcohol frontloading [60]. In general, this approach uses a change point detection algorithm [67] to find changes in the drinking rate using the cumulative consumption throughout the session. If these changes in drinking rate indicated that during the first half of the session the animal drank (1) pharmacologically relevant quantities of alcohol at (2) a faster rate than later in the session, the session was categorized as a frontloading session. If the changes in the drinking rate were not well classified using the change point detection algorithm, the drinking pattern was classified as inconclusive. See [60] for full details and example software.

### 2.4 Head-Fixed Apparatus

We used a modified version of the International Brain Laboratory open-source head-fixed mouse behavior apparatus [61] for all head-fixed experiments (Figure 1 B). A detailed description of the head-fixed apparatus used by IBL is available at [68] and we supply a description of the modifications we used in [69], including a detailed description of the fluid delivery system with a parts list. We recorded video using monochromatic Chameleon 3 cameras (Teledyne Flir) at approximately 30 Hz. The cameras were modified with IR filters to only allow IR light to be detected and the animal was illuminated using an IR spotlight. We ran the task using a Bpod interface box (Sanworks) that was controlled using custom Matlab (Mathworks) software. An IR LED was used to synchronize video frames with task events. A rubber wheel (Lego) was positioned below the animal’s front paws, but it was locked in place. A video screen was positioned in front of the animal, but no visual stimuli were provided. Rather, the screen displayed a gray background throughout the session.

For cohort 1, an IR lickometer (Sanworks) was used to measure licks by detecting beam breaks near the tip of the sipper (Figure 1 C). For cohort 2, the IR lickometer was removed and replaced with a real-time machine vision lick detection system that used an OpenMV4 CAM H7 camera (OpenMV) (Figure 1 D). Example videos were gathered from cohort 1 mice after the completion of their experiment. Individual image frames were subtracted from a non-licking frame to highlight jaw and tongue movements associated with licking and cropped to reduce image size and improve performance. These training frames were hand scored and used to train a deep net using tensorflow to recognize licks. This network was then installed on the camera and run in real time to detect licks in cohort 2.

Animal body parts were tracked using DeepLabCut [45]. In cohort 1, the animal’s left paw and snout tip were tracked. In cohort 2, the animal’s left paw, snout tip, and lower jaw were tracked, as well as the sipper tip. Sipper location and lower jaw position were used to refine lick detection post-hoc. In general, licks were registered when the animal’s jaw moved below the sipper tip.

Fluid delivery was controlled by a Pump 11 Elite syringe pump (Harvard Apparatus) that was triggered using the custom Matlab (Mathworks) software used to run the Bpod controller. Fluid was delivered to the animal using a custom stainless steel sipper tube. This tube contained a concentric smaller tube that was connected to a vacuum line via a computer controlled pinch valve and a collection tube. This system allowed for fluid to be dispensed onto the end of the sipper and for unconsumed fluid to be suctioned off and collected following the end of the trial.

### 2.5 Head-Fixed Acclimation and Task

Mice were acclimated to head-fixation using a two-step process. On the first day, approximately 1 hour in the mice’s dark cycle, each animal was placed on the mouse holder, but not head-fixed. The mouse holder was removed from the head-fixation system and placed in a large bin outside the enclosure chamber that holds the apparatus. The mouse was allowed to explore the holder for 10 minutes and then was returned to its home-cage in the vivarium for approximately 2 hours. Then, the mouse was retrieved and head-fixed in the mouse holder for 10 minutes outside the apparatus. No fluids were available during this period.

On the next day, the mice were head-fixed in the full head-fixation apparatus and put through the standard Pavlovian task utilized throughout the experiment (Figure 1 E). In this task, a 5 kHz tone played for 1 second, then a 4 μL droplet of fluid was dispensed onto the end of the sipper. If the animal did not lick for 15 seconds, the droplet was suctioned off the sipper. If the animal licked during the access period, the animal would have 5 more seconds to consume the droplet before any remaining fluid would be suctioned off the sipper. The session lasted 30 minutes. During the two-week acclimation period, the fluid was exclusively 20% (v/v) ethanol (Decon Laboratories, Inc.) and mice were put through the task at various times between 1 and 3.5 hours into their dark cycle.

On the Monday of the second week of head-fixed drinking acclimation (day 5 of head-fixed drinking), 50 µL of retro-orbital blood was drawn immediately after the head-fixed session. Blood samples were immediately centrifuged, and plasma was withdrawn and stored at -20°C. Blood ethanol concentration (BEC) measurements were performed one week later using an AM1 alcohol analyser (Analox Instruments).

### 2.6 Head-fixed Quinine Tests

Following 2 weeks of head-fixed acclimation drinking, the mice underwent two weeks of identical aversion-resistant drinking assessments where increasing concentrations of quinine were introduced into the ethanol (Figure 1 A). Each week, the concentration of quinine increased from 0 g/L up to 3.6 g/L. Two different sets of quinine concentrations were used in the two cohorts to probe a larger dynamic range of quinine consumption.

### 2.7 Home-Cage Quinine Tests

After head-fixed drinking, the mice underwent a quinine drinking test in home-cage to compare head-fixed ARD to home-cage ARD (Figure 1 A). In these tests, mice received access to 20% ethanol (v/v) (Decon Laboratories, Inc.) and no water on Monday through Wednesday. Access was provided approximately 1.5 hours into the dark cycle (similar time of day used for head-fixed drinking) for 1 hour. On Thursday, 0.9 g/L quinine was added to the alcohol. On Friday, a higher concentration of quinine was used (3.6 g/L for cohort 1, 1.8 g/L for cohort 2). Alcohol and quinine-adulterated alcohol was made available to the mice in sippers constructed of 10 mL graduated pipettes (readable to +/- 0.1 mL) outfitted with ball-bearing sippers. The volumetric drinking monitor system was not used in order to prevent contamination of the system with quinine.

### 2.8 Body Part Movement Analysis

Snout and paw positions were found for each video frame using DeepLabCut [45]. Low confidence coordinates (less than 0.5) were linearly interpolated with the temporally nearest high confidence coordinates. Snout and paw speeds were calculated as the change in 2-dimensional pixel position divided by the time between video frames. Five sessions were excluded from correlation analyses with body part speeds due to substantially higher or lower consumption values that produced large relative consumption outliers that biased correlation calculations.

### 2.9 Data and Analysis Software

Raw data were first processed and analyzed in Matlab (Mathworks). Processed data were subjected to statistical assessments and plotted in Prism (GraphPad). ANOVAs (including repeated measures), t-tests, other tests were used and corrections (e.g., Greenhouse–Geisser) were applied when appropriate. When appropriate, sex was considered as a factor in ANOVAs. For the sake of brevity, main effects and interactions that were not found to be significant were omitted from discussion. All organized data, analysis software, behavioral code, machine vision code for OpenMV cameras, plotting files, and a detailed description of the head-fixed apparatus are freely available [69]. Raw data and video files are available upon request due to their size.

## 3 Results

### 3.1 Drinking in the Dark

After implantation of head-fixation bars and recovery from surgery, mice underwent drinking in the dark (DID) for 1 week (5 weekdays) (Figure 2). In terms of consumption throughout the week, no main effects were observed based on day, sex, or cohort, though there was a sex by day interaction (F(4,73) = 2.501, p = 0.0499) (Figure 2 A). We assessed alcohol frontloading – an alcohol drinking pattern where intake is skewed toward the onset of access and which results in intoxication – using statistical methods to identify changes in drinking rate (Figure 2 B) [60]. This method sought to identify drinking patterns that result in high consumption rates that outpaced the rate of alcohol metabolism near the start of a drinking session. Mice that engaged in frontloading rapidly experienced the pharmacological effects of alcohol early in the session because they could not metabolize the alcohol fast enough to keep up with consumption. We observed distinct patterns of frontloading throughout DID (Figure 2 C). Sessions that were categorized as inconclusive with regards to frontloading using the statistical analysis were recategorized by comparing the cumulative consumption through the first 15 minutes to the average cumulative consumption traces for well classified patterns on each day. We categorized mice as frontloaders or non-frontloaders based on their drinking pattern on day 5 of DID, unless all previous days were the opposite type, in which case we used the category from the previous days. On the last day of DID, frontloading patterns were significantly higher during the initial portion of the session compared to non-frontloading patterns (Figure 2 D, FDR controlled rank sum test p < 0.01). No difference was found in session-wide consumption based on frontloader status (Figure 2 E, t(21) = 0.066, p = 0.95). Although females appeared to have a higher proportion of non-frontloaders compared to males, this difference was not significant (Figure 2 F, Fisher’s exact test, p = 0.371). Also, no significant differences were observed in the proportion of mice identified as frontloaders between cohorts (Cohort 1: 3/11, Cohort 2: 4/12, Figure 2 F, Fisher’s exact test, p > 0.99).

**Figure 2:**
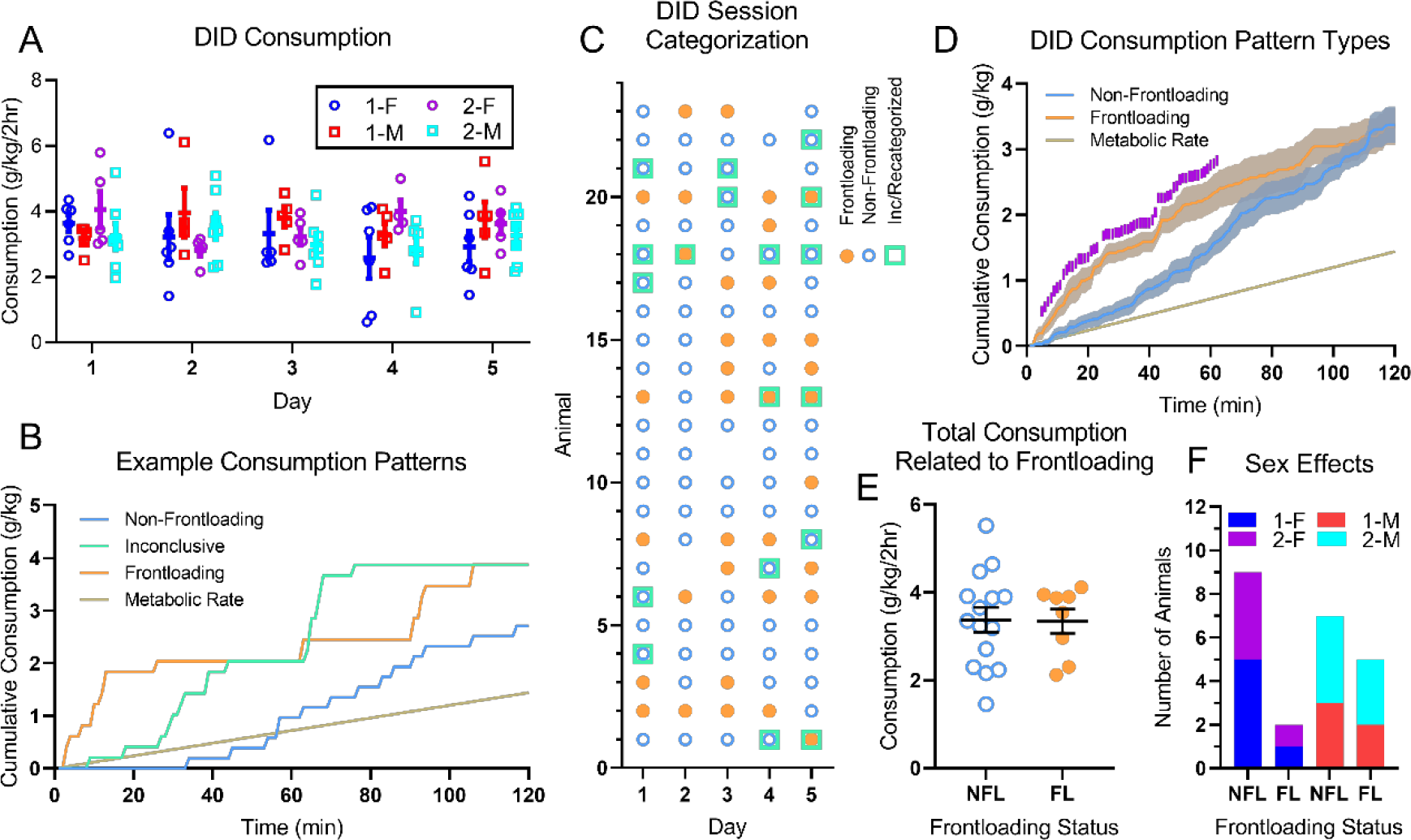
Drinking in the dark consumption patterns differentiate frontloaders and non-frontloaders. **(A)** Consumption levels were stable throughout DID and no main effects of sex or cohort were found. Cohort numbers (1,2) and sex (M,F) are listed in the legend. **(B)** Example cumulative consumption graphs for three different DID sessions that clearly demonstrate frontloading, non-frontloading, and inconclusive drinking patterns in relation to metabolic rate for alcohol [70]. Consumption levels above the metabolic rate indicate that the animal experienced the pharmacological effects of alcohol because it was not able to metabolize alcohol fast enough to outpace consumption [70]. **(C)** Frontloading classification for each drinking session. Sessions with missing data omitted. **(D)** Frontloading and non-frontloading average cumulative consumption on the last day of DID. Frontloading sessions tended to have higher drinking during the initial hour of the session (FDR controlled rank sum test p < 0.01 for each time bin marked with purple tick). **(E)** No differences were observed between frontloading and non-frontloading sessions on the last day of DID in terms of overall consumption on the last day of DID. **(F)** Though females had a higher proportion of non-frontloaders, no significant effects of sex were observed in terms of frontloader status representation. Also, no significant effects of cohort were found on frontloader status representation.

### 3.2 Head-Fixed Alcohol Acclimation

Mice were acclimated to head-fixation on a Monday with a 10-minute exploration of the head-fixed apparatus in the morning, followed by 10 minutes of head-fixation. During the following 9 weekdays, the mice were head-fixed and participated in the task (Figure 3). Mice generally consumed 20% alcohol in the task (Figure 3 A). Cohort 2 drank significantly more, drinking decreased slightly over days, and one interaction was observed (main effect of cohort: F(1,19) = 53.22, p < 10^-4^, main effect of day: F(4.773,88.91) = 2.935, p = 0.018, day by cohort interaction: F(8,149) = 2.25, p = 0.027, no effects of sex). We believe increased consumption in cohort 2 was due to better sipper alignment with the mice’s mouths due to the removal of the IR lickometer and improved visibility of the sipper.

**Figure 3:**
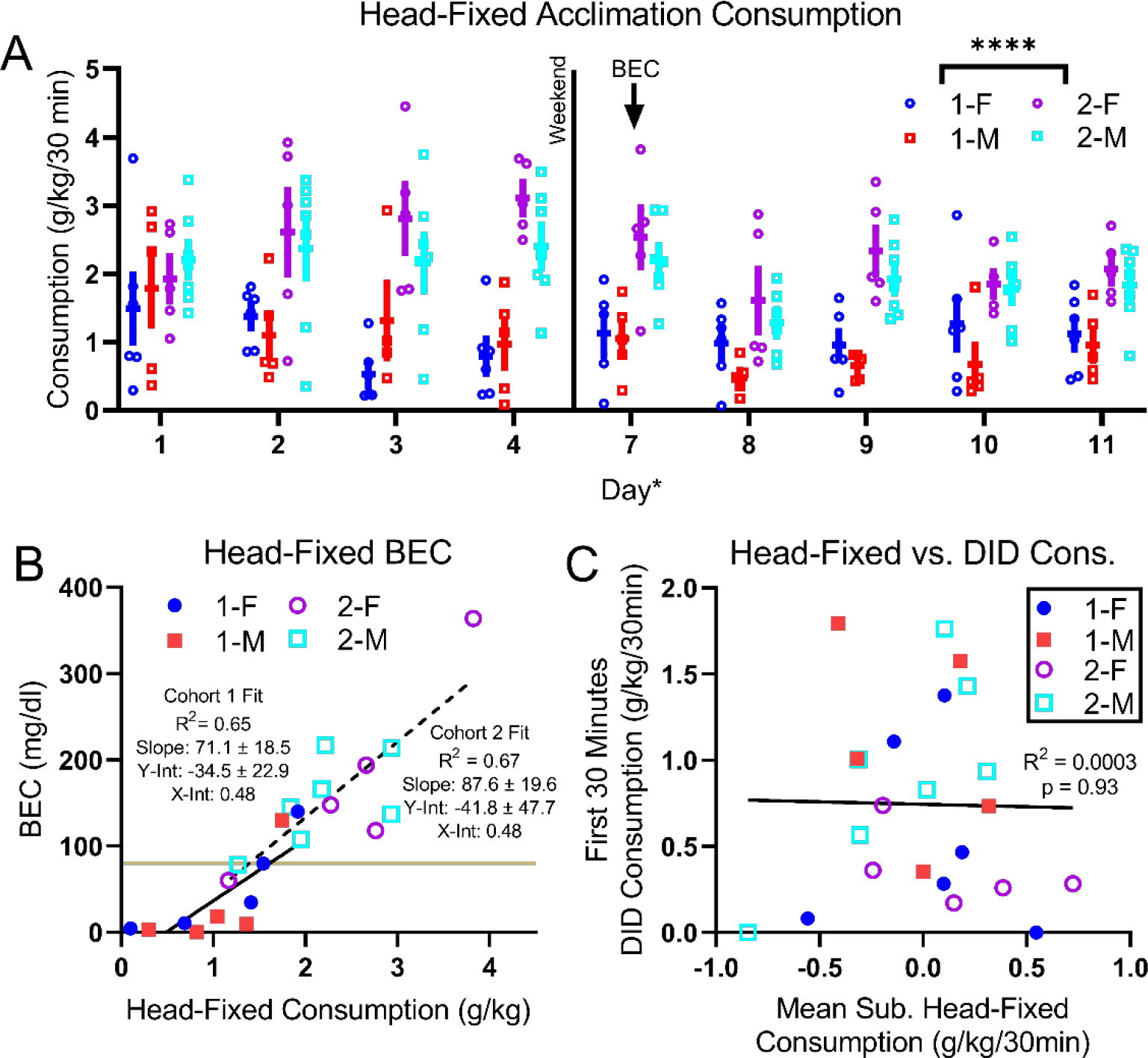
Head-fixed mice readily consumed 20% alcohol. After DID sessions, mice received a 10-minute exploration period of the head-fixed apparatus and then a 10-minute head-fixed acclimation with no alcohol (day 0, no data). Mice then received nine 30-minute sessions with the task and 20% ethanol. **(A)** Alcohol consumption through acclimation. Cohort 2 mice drank significantly more than cohort 1 and drinking changed significantly over days (^*^: p < 0.05, ^****^: p < 10^-4^). **(B)** Blood ethanol concentrations (day 7 (A)) correlated with consumption levels in both cohorts and the fits from each cohort agreed within error. Many mice exhibited pharmacologically relevant BEC values (greater than 80 mg/dl, brown line). **(C)** Head-fixed consumption was unrelated to average DID consumption during the first 30 minutes of the DID session. Head-fixed consumption was cohort mean subtracted to control for increased consumption in cohort 2.

BECs were measured on day 7 (Monday) and were well correlated with consumption values (Figure 3 B). Fits were assessed for each cohort separately due to the main effect of cohort on consumption. Fits from both cohorts agreed within error. Head-fixed consumption was unrelated to DID consumption during the first 30 minutes of the DID session (Figure 3 C). We chose to use the first 30 minutes of the DID session for comparison because the head-fixed drinking sessions were 30 minutes in duration. The mean head-fixed alcohol consumption (1.62 g/kg/30 minutes) was more than double the mean DID session consumption during the first 30 minutes (0.74 g/kg/30 minutes).

### 3.3 Licking and Body Movement Behavior

We examined licking behavior and movement of the snout tip and left paw during the head-fixed task (Figure 4). We focused on the alcohol baseline day for the first quinine test week to maximize acclimation to alcohol drinking in the head-fixed task. Lick rate time locked to the tone demonstrated that mice tended to lick immediately following sipper clear (the moment suction was applied to remove any remaining drop from the sipper tip) and after the drop had been extruded onto the sipper tip (Figure 4 A). When we averaged the lick rate during epochs of interest, we found that lick rate varied by epoch and we found an interaction between epoch and sex, but no effect of cohort (Figure 4 B, main effect of epoch: F(2.201,41.8) = 18.9, p < 10^-4^, epoch by sex interaction: F(4,76) = 3.17, p = 0.018). We then compared the pre-tone epoch lick rate to the lick rates during the other epochs of interest (Dunnett’s multiple comparisons test) and found significant increases during the sipper clear, drop, and the post-drop epochs. Next, we examined the snout tip and paw speeds throughout the trials. Individual trial speeds tended to be somewhat noisy with snout speeds showing a higher baseline level, but paw speeds being marked by large deviations caused by rapid movements of the paw (Figure 4 C). Both snout tip and paw speeds increased following the sipper clear time point but showed few changes during tone or drop time points (Figure 4 D). Both snout tip (Figure 4 E) and paw (Figure 4 F) speeds increased before and during licking.

**Figure 4:**
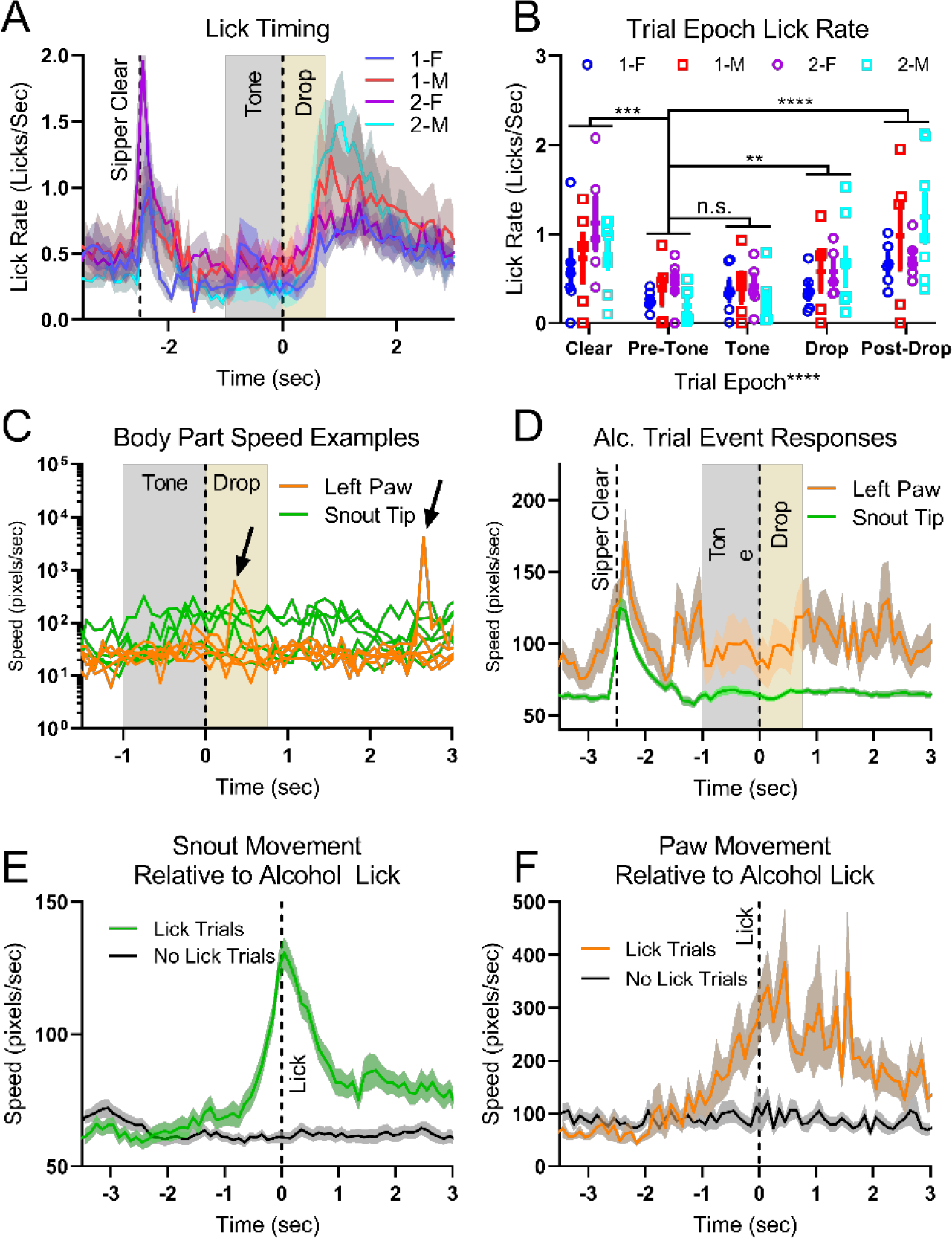
Body part movement was related to alcohol drinking behaviors. **(A)** Average lick rate time locked to the tone (mean +/- SEM across mice, each mouse averaged across all trials on quinine test week alcohol only baseline day 1). The drop time period refers to the period during which the drop was extruded on the end of the sipper. **(B)** Lick rates from (A) were averaged in epochs of interest (Clear: -2.5 to -2 sec, Pre-Tone: -1.5 to -1 sec, Tone: -1 to 0, Drop: 0 to 0.75 sec, Post-Drop: 0.75 to 1.75). **(C)** Five example body part speed profiles time locked to the tone (first five trials for animal 1). Note that the snout tip speed tended to be higher than the paw speed at baseline, but that the paw speed exhibited large spikes in speed (arrows). **(D)** Body part speed profiles (mean +/- SEM across mice, each animal averaged across all trials during the session). Both profiles exhibited peaks near the sipper clear time, but few other changes related to the tone or drop release. The paw speed was generally higher and more variable compared to the snout speed. **(E)** Snout speed time locked to the first lick on a trial and similar time points from trials without licks. The snout speed increased during and immediately before the lick. **(F)** Paw speed time locked to the first lick on a trial and similar time points from trials without licks. The paw speed increased immediately before and during the lick. (^*^: p < 0.05, ^**^: p < 0.01, ^***^: p < 0.001, ^****^: p < 10^-4^.)

### 3.4 Aversion-Resistant Drinking

Following acclimation to head-fixed alcohol drinking, mice underwent a sequence of two quinine test weeks wherein quinine concentrations increased throughout the week (Figure 5)._We adjusted the second cohort’s quinine concentrations downward to probe a wider range of consummatory responses. Comparing consumption values for only quinine concentrations that were tested in both cohorts (0, 0.3, 0.9, and 1.8 g/L), cohort 2 drank more and increasing quinine concentration reduced consumption, as expected (Figure 5 A, main effect of quinine concentration: F(4.48,85.18) = 7.06, p < 10^-4^, main effect of cohort: F(1,19) = 72.34, p < 10^-4^, and quinine concentration by cohort interaction: F(7,133) = 2.55, p = 0.017, no effect of sex). The increased consumption in cohort 2 may have been driven by improvements to sipper alignment, as well as the slower increase in quinine concentration used in cohort 2 during the initial days of quinine exposure which would allow for easier habituation to the taste of quinine. We examined the stability of the drinking response across the two quinine test weeks by comparing consumption for each animal at a given quinine concentration (Figure 5 B). Though we found cohort 2 mice drank more than cohort 1 overall and we found a cohort by week interaction, we found no main effect of week, so drinking levels did not change during the second quinine test (main effect of cohort: F(1,88) = 71.37, p < 10^-4^, week by cohort interaction: F(1,88) = 4.28, p = 0.042, no effect of sex). When we directly related the first quinine exposure consumption to the second quinine exposure consumption for each animal, we found the two values were only weakly correlated (Figure 5 B). This instability in quinine test-retest results has been observed previously in home-cage drinking (Anna Radke, personal communication). Based on this instability, for the remainder of the head-fixed quinine analysis, we only considered the first week of quinine exposure.

**Figure 5:**
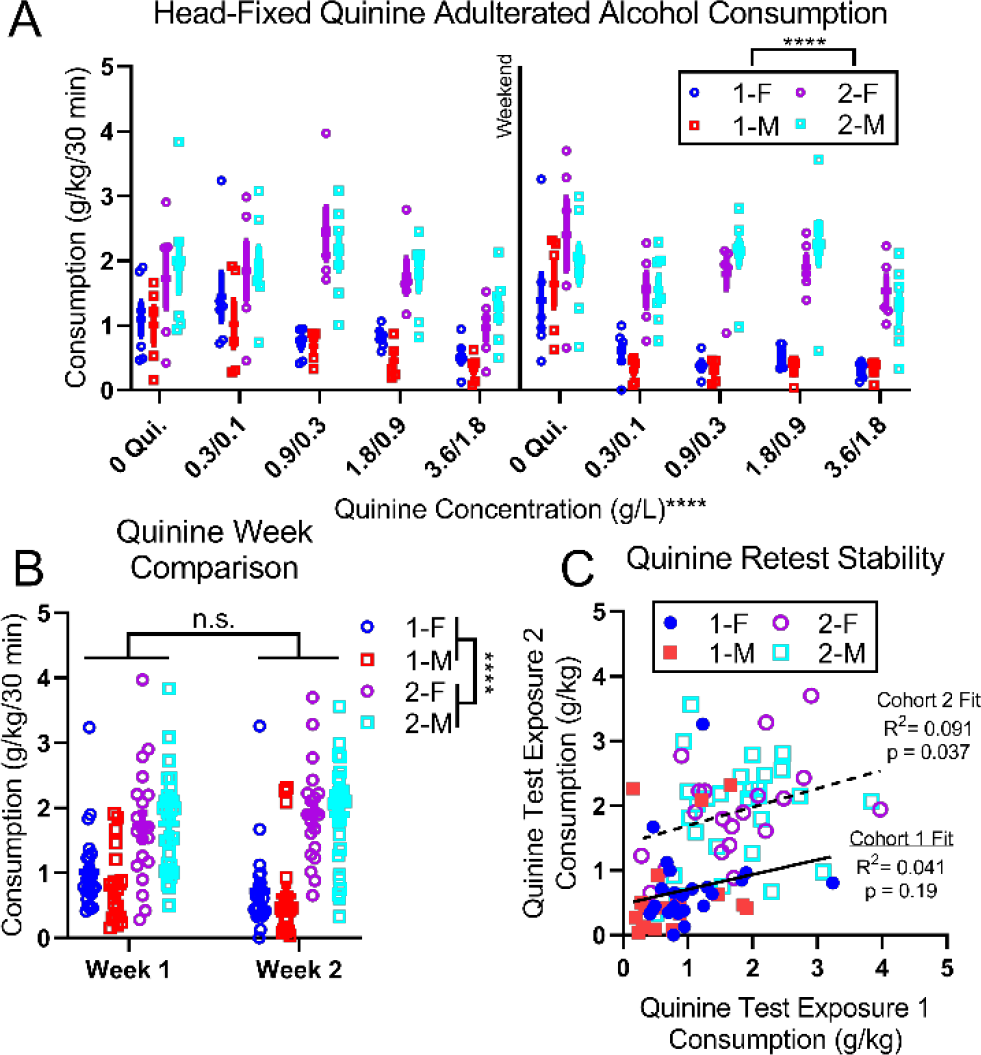
Head-fixed mice readily consumed quinine adulterated alcohol. **(A)** Consumption decreased with quinine concentration during two quinine test weeks, though more robustly for cohort 1. Cohort 1 received quinine concentrations of 0.3, 0.9, 1.8, and 3.6 g/L. Cohort 2 received quinine concentrations of 0.1, 0.3, 0.9, and 1.8 g/L. **(B)** Comparison of consumption values for matched quinine concentrations. Cohort 2 drank more in general, but drinking did not change during the second exposure. **(C)** Consumption for matching quinine concentrations across the two test weeks was uncorrelated for cohort 1 and weakly correlated for cohort 2. In B and C, only quinine concentrations used in both cohorts (0, 0.3, 0.9, and 1.8 g/L) were considered.

We examined licking behavior and consumption for the alcohol baseline day and the 0.9 g/L quinine days during the first week of quinine exposure (Figure 6 A-D). Mice that were classified as frontloaders during DID were found to lick more during the alcohol baseline day (Figure 6 A) and the 0.9 g/L quinine day (Figure 6 C). Though DID frontloaders had similar overall DID consumption values over the two hour DID session as compared to non-frontloaders (Figure 2 E), DID frontloaders drank more alcohol in the 30 minute head-fixed drinking session (Figure 6 B, t(21) = 2.11, p = 0.047). Given the shortness and paced nature of the head-fixed session, we did not assess frontloading in the head-fixed session. DID frontloaders did not consume more alcohol adulterated with 0.9 g/L quinine, though DID non-frontloaders drank less than the cohort mean subtracted alcohol baseline day (one sample t-test comparison to 0, t(15) = 2.648, p = 0.018).

**Figure 6:**
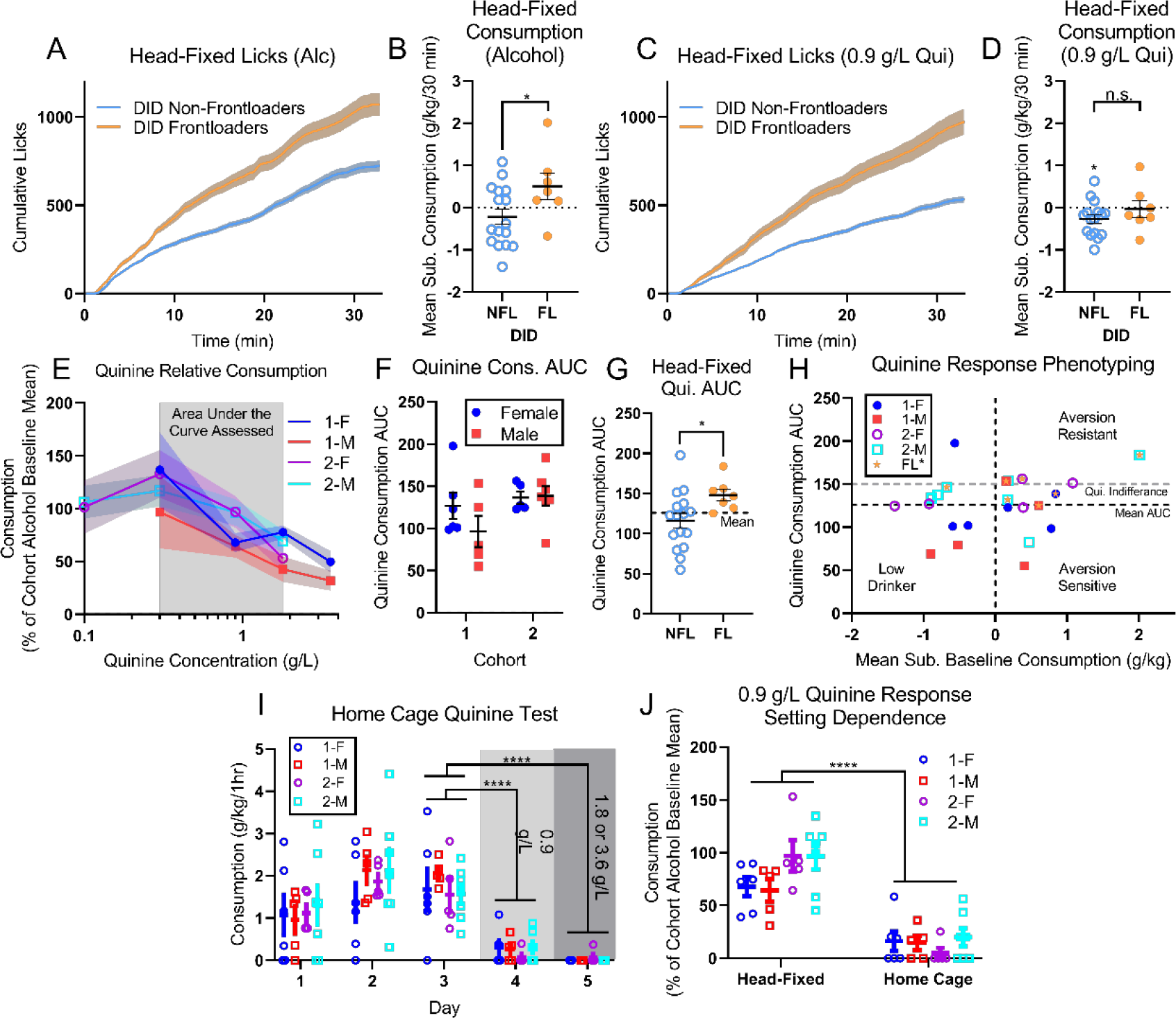
Head-fixed mice exhibited aversion-resistant alcohol drinking for quinine. **(A)** DID frontloaders licked more throughout the head-fixed alcohol session of the first quinine test week than non-frontloaders and **(B)** consumed more alcohol (cohort mean consumption subtracted to account for cohort effects in consumption). **(C)** On the 0.9 g/L quinine day of the first quinine test week, DID frontloaders also licked more. **(D)** DID frontloaders did not drink more in this session but DID non-frontloaders drank significantly less alcohol than the alcohol baseline day. **(E)** Consumption levels for the first week of quinine testing (Figure 5 A) relative to the cohort average alcohol consumption baselines on the first day of the week (mean +/- SEM across mice). **(F)** Area under the curve (AUC) was calculated for overlapping quinine concentrations from E. No differences were observed based on cohort or sex. **(G)** Quinine consumption AUC for each DID frontloading type. DID frontloaders had elevated quinine consumption AUC. **(H)** Animal consumption (cohort mean subtracted) vs. quinine consumption AUC. Mice could be phenotyped by comparing AUC to alcohol consumption. Gray line corresponds to AUC with average at 100% in A (i.e., indifference to quinine). DID frontloaders (orange star) tended to be aversion-resistant. **(I)** Consumption in a follow-up home-cage drinking test showed decreased consumption for two concentrations of quinine. **(J)** Relative consumption during quinine adulteration in head-fixture was higher than in home-cage for the common concentration of 0.9 g/L. (^*^: p < 0.05, ****: p < 10^-4^)

We compared the consumption of each animal during the first week of quinine exposure to the cohort mean alcohol only consumption (Figure 6 E). As expected, consumption tended to decrease as quinine concentration increased (main effect of quinine over common concentrations (0.3-1.8 g/L): F(2,38) = 12.32, p < 10^-4^, no effect of sex). To quantify quinine consumption in a single measure, we calculated the area under the curve (AUC) for the common quinine concentrations (0.3-1.8 g/L, Figure 6 F). We found no effects of sex or cohort on quinine consumption AUC when examining relative consumption, despite the elevated alcohol and quinine adulterated alcohol consumption levels we observed in cohort 2. DID frontloaders had higher head-fixed quinine consumption AUC than non-frontloaders (t(21) = 2.18, p = 0.041, Figure 6 G). We used alcohol consumption (cohort mean subtracted) and quinine consumption AUC to phenotype the mice (Figure 6 H) as aversion-resistant (above cohort mean alcohol consumption and above mean quinine consumption AUC), aversion-sensitive (above cohort mean alcohol consumption and below mean quinine consumption AUC), and low drinkers (below cohort mean alcohol consumption and below mean quinine consumption AUC). Of 23 mice included in the study, 7 were classified as aversion-resistant and 3 others produced measures that were very close to aversion-resistance (above cohort mean alcohol consumption and slightly below mean quinine consumption AUC). Of these 10 mice, 6 were classified as frontloaders during DID, indicating that DID frontloaders were more likely to exhibit aversion-resistant drinking in head-fixture (Fisher’s exact test, p = 0.019). No effects of sex (Fisher’s exact test, p > 0.99) or cohort (Fisher’s exact test, p = 0.68) were observed. Quinine consumption AUC and aversion-resistance classification taken together indicate that mice that exhibited frontloading during DID also exhibited more aversion-resistant drinking in head-fixation.

Following head-fixed quinine testing, mice were exposed to quinine in their home-cages (Figure 6 I). Mice were given one hour of access to only alcohol for three days (M-W). On the fourth and fifth day, quinine was added to the alcohol in increasing concentrations. Consumption decreased with the addition of quinine, as expected (main effect of day F(2.905,55.19) = 40.15, p < 10^-4^, Dunnett’s multiple comparisons test between day 3 and days 4 and 5, p<10^-4^, no effect of sex). The relative consumption of alcohol adulterated with 0.9 g/L quinine was higher when mice were head-fixed than when in home-cage (Figure 6 J, main effect of setting: F(1,19) = 97.06, p < 10^-4^, setting by cohort interaction: F(1,19) = 5.98, p = 0.024, no effect of sex).

### 3.5 Aversion-Related Body Movement

Clear behavioral responses to quinine were observed in the animal behavior (e.g., rapid paw movement and twitching). We focused on comparisons between alcohol, low quinine (0.1 g/L), and high quinine (3.6 g/L) to capture the most robust behavioral responses. We quantified these behaviors by analyzing body part tracking from DeepLabCut of the animal’s left paw and snout tip (Figure 7). For snout tip, we saw elevated speeds relative to alcohol baseline, especially immediately after licking (Figure 7 A and B). Interestingly, we also saw depressed speeds on the lowest quinine concentration near the same time points. The snout tip speed immediately following licking was inversely related to quinine consumption relative to baseline (Figure 7 C). In other words, the less the animal drank, the more it moved its nose when it drank. For the paw, we saw elevated speeds at the highest quinine concentration throughout the trial and especially immediately after licking (Figure 7 D and E). The paw speed immediately following licking was inversely related to the quinine consumption relative to baseline in Cohort 1, but not Cohort 2 (Figure 7 F). In other words, like with snout speed, the less the animal drank, the more it tended to move its paw when it drank.

**Figure 7:**
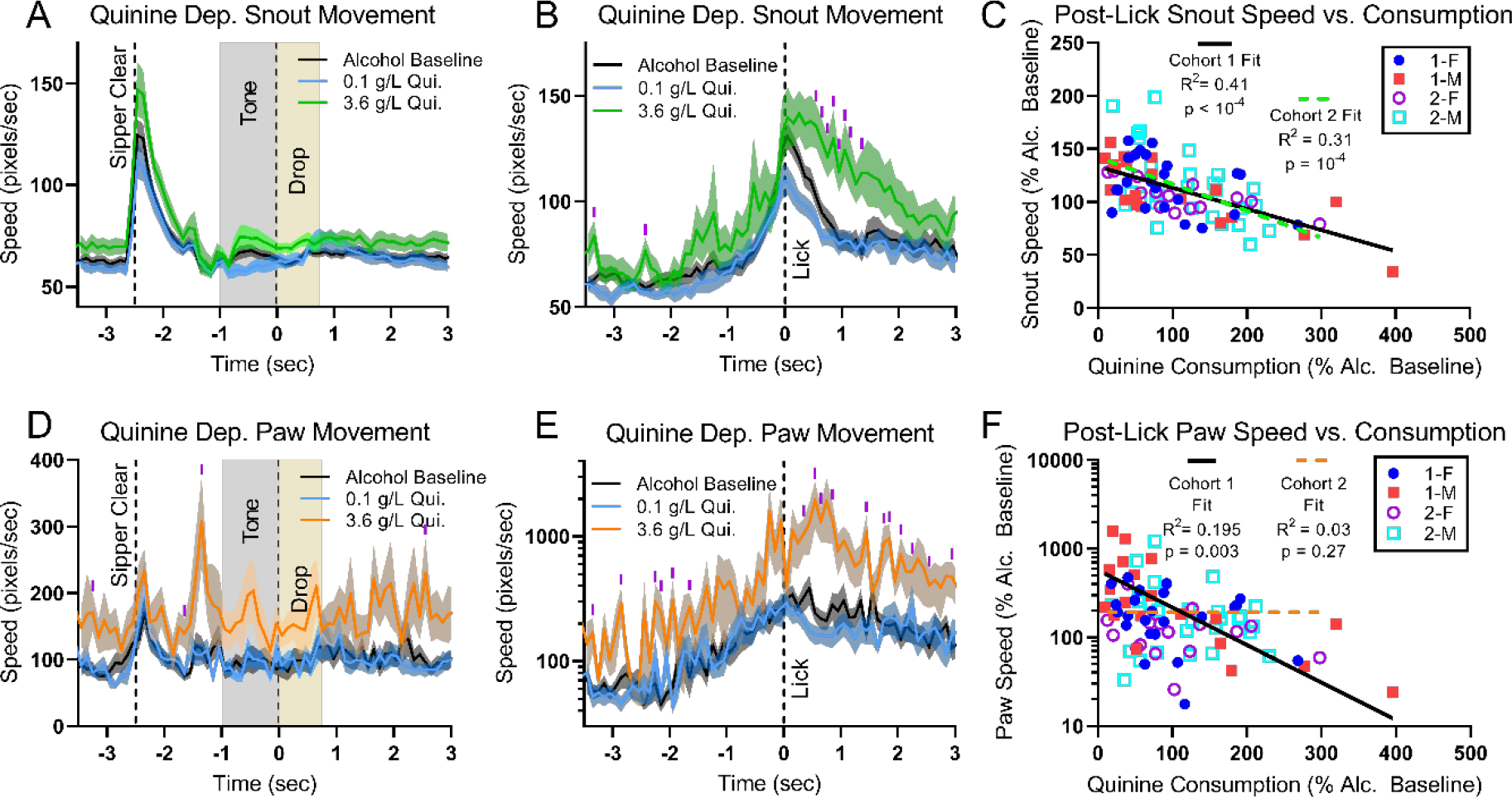
Body part movement was related to aversion-resistant drinking. Snout speed time locked to the beginning of each trial **(A)** and the first lick of the trial **(B)** increased immediately after licking. **(C)** The snout speed was inversely related to the relative consumption of quinine across all first week quinine sessions. Paw speed time locked to the beginning of each trial **(D)** and the first lick of the trial **(E)** (note log scale) increased for the highest quinine concentration. **(F)** The paw speed was inversely related to the relative consumption of quinine across all first week quinine sessions for Cohort 1, but not Cohort 2 (semi-log fit, F-test against slope equal to 0). (A, B, D, E: Mean +/- SEM across mice shown, FDR controlled t-test p < 0.05 for each time bin marked with purple tick).

## 4 Discussion

### 4.1 Alcohol Consumption and Aversion-Resistant Drinking during Head-Fixation

In this study, we found that head-fixed C57BL/6J female and male mice readily consumed alcohol to the point of intoxication and that this population of head-fixed mice exhibited heterogenous levels of aversion-resistant drinking. We showed that this was possible with relatively short drinking histories (1 week of DID). We also found that mice consumed higher concentrations of quinine when head-fixed than when in home-cage.

A key concern for the application of head-fixed methods to studies of alcohol drinking is the role of restraint stress associated with head-fixation, especially given the well-established relationship between stress and alcohol addiction [71, 72]. Fortunately, while it has been shown that the initial days of head-fixation produce a stress response, mice habituate to head-restraint and stress levels return to normal [73]. Furthermore, no differences have been reported in alcohol self-administration following restraint stress in a genetically diverse mouse strain [74]. Regardless, in the future, it will be critical to monitor corticosterone levels in mice during quinine consumption to better understand the impact of stress in our experimental paradigm.

In relation to concerns about stress, we observed that mice drank higher concentrations of quinine in head-fixture than when in home-cage. It is possible that these results are due to stress, but we believe this is unlikely due to habituation to head-fixation and our observations of the mice. We believe elevated quinine consumption in head-fixture is due to the lack of behavioral freedom (i.e., there is nothing else to do but lick or not lick) in comparison to home-cage. We wish to emphasize that the reduction in behavioral freedom in head-fixation relative to home-cage still permits the mice to abstain and many mice reduced their consumption levels substantially for quinine adulterated alcohol. In other words, though head-fixation may make it easier to engage in higher levels of ARD relative to home-cage, it still permits aversion-resistant and aversion-sensitive drinking behavior. Thus, we believe head-fixation is a self-administration paradigm in which ARD can be studied.

A similar concern is whether the high levels of consumption that we observed in head-fixation – roughly double in head-fixation (1.6 g/kg/30 minutes) vs. home-cage DID (0.75 g/kg/30 minutes) – is real and, if so, fueled by stress. Based on the strong correlations we observed between consumption values and BEC (Figure 3 B), we believe the high drinking levels we observed for head-fixation were the result of consumption. As with elevated quinine consumption relative to home-cage, we believe this is due to lack of behavioral freedom during head-fixation, not stress. Also, note that head-fixed consumption was more limited, both in terms of overall time (30 minutes in head-fixture vs. 2 hours in DID) and the pacing (paced trials in head-fixture vs. unlimited self-administration in DID). It is known that limiting access to alcohol can increase alcohol consumption in rodents [75], so the limited access structure of head-fixation may have also increased consumption of alcohol and quinine adulterated alcohol.

We also observed significant differences between cohort 1 and 2 in terms of alcohol and quinine adulterated alcohol consumption that we believe were caused by the switch from an IR lickometer to measure licks in cohort 1 to a real-time machine vision system to measure licks in cohort 2. This change allowed us to better position the sipper near the mouth of the animal because the animal’s mouth was not obscured by the IR lickometer. We believe this change facilitated easier access to the fluid and increased drinking. It is also possible that the slower increase in quinine concentration used in cohort 2 eased habituation to quinine and increased consumption levels, but we do not believe this was a major factor given the lack of a cohort effect in relative quinine consumption. Though we utilized a sipper alignment procedure to maintain consistent positioning of the sipper for each animal across days, in the future we plan to develop improvements to this sipper alignment procedure to better unify the sipper location across mice.

### 4.2 Frontloading

It has been argued that alcohol frontloading – a skew in drinking towards the beginning of a drinking session – is driven by the rewarding effects of alcohol and can be used as a measure of the motivation to consume alcohol [60]. We quantified frontloading behavior during home-cage DID sessions and related the frontloader status of each animal to head-fixed consumption. We found that in head-fixation, DID frontloaders licked more, consumed more alcohol, and were more aversion-resistant. We believe these similarities in results across home-cage DID and head-fixation reflect enhanced motivation for alcohol in a subset of the mice and that head-fixation is a viable paradigm for modeling AUD.

Though we compared DID frontloading to head-fixed ARD, we did not compare DID frontloading to home-cage ARD. We did assess consumption of quinine adulterated alcohol in home-cage for comparison to head-fixed consumption of quinine adulterated alcohol, but the concentrations of quinine used in these home-cage tests were too high to effectively assess ARD in home-cage. Several other studies have assessed DID frontloading and quinine consumption in home-cage [13, 21, 76], but we are not aware of any studies that directly report on the relationship between DID frontloading and future ARD in home-cage. In the future, it would be helpful to establish whether DID frontloading predicts future ARD in home-cage.

### 4.3 Non-consummatory behaviors as an indication of ARD

We utilized machine vision techniques to track the left paw and snout movements of the mice during head-fixation [45]. Using these data, we showed that it was possible to relate behaviors other than drinking (i.e., paw and snout movement) to the aversiveness of the quinine (e.g., snout movement and quinine consumption were inversely correlated). This relationship between aversiveness and movement was readily apparent from visual observation of the mice, which corresponds well with previous reports that outbred mice respond to oral quinine administration with aversive oral-facial response [77]. In the future, we hope to utilize more advanced techniques to categorize mouse movements (e.g., [54, 55]) beyond the simple speed measurements we utilized. Developing better behavioral analyses [78] and relating them to neural data offer the potential to substantially improve our understanding of complex behaviors like ARD.

### 4.4 Aversion-Resistant Drinking and Compulsivity

The relationship between aversion-resistant drinking and so-called compulsive drinking has been a source of discussion in the AUD research field, with some authors challenging the view that aversion-resistant drinking can be equated with compulsive drinking in pre-clinical animal models in particular [3]. Prior work (including ours [18, 19]) has equated compulsive drinking with aversion-resistant drinking. Aversion-resistant drinking can be directly assessed behaviorally by investigating the degree to which an animal will continue to consume alcohol despite negative consequences such as quinine or footshock. In contrast, compulsivity is more difficult to define, but crucially involves a feeling that motivates someone to pursue some action that is “not in line with one’s overall goal” [79]. It is possible for various factors to contribute to aversion-resistant drinking, including factors that are unrelated to the subjective motivation to drink, such as differing sensitivity to the aversive stimulus [3]. However, it is difficult for us to understand how an animal could engage in aversion-resistant drinking, but completely lack an internal motivation for alcohol necessary for compulsive drinking. The current data, which indicate that frontloaders are more willing to consume higher concentrations of quinine-adulterated alcohol, suggest that subjects with a higher motivation for alcohol consumption may be more likely to engage in aversion-resistant drinking. Still, our inability to conceptualize an alternative explanation is not direct evidence for the existence of this subjective feeling on the part of the animal. Therefore, we believe these two phrases should be delineated and the appropriate term for non-human studies is usually aversion-resistant drinking. We agree with [3] that this subjective feeling is central to compulsivity and that it is unclear how this feeling could be assessed in non-humans where it can be extremely difficult to detect and characterize their subjective experiences.

Fortunately for pre-clinical research, insofar as the subjective motivational drive to drink that is associated with compulsive drinking is concerned, compulsive drinking is less heavily weighted than aversion-resistant drinking in the DSM-V symptoms that are used to assess AUD in humans [1]. Only one symptom (“In the past year, have you wanted a drink so badly you couldn’t think of anything else?”) addresses the subjective motivational drive to drink, whereas six of the symptoms are directly related to continuing to drink despite negative consequences (e.g., “In the past year, have you continued to drink even though it was causing trouble with your family or friends?”). Therefore, we believe investigating aversion-resistant drinking in pre-clinical animal models is appropriate and the most tractable use of these model systems. The precise details of these studies, such as the type of aversive stimulus and the drinking history of the animals will affect the conclusions that should be drawn from these animal systems. For instance, drinking history may affect the neurobiological mechanisms that underly ARD such that results from longer drinking history animals may be more relevant to patients suffering from a long history of AUD, whereas results from shorter drinking history animals may be more relevant to patients near the start of AUD.

### 4.5 Future Directions

In the future, we will use high-density electrophysiology [37, 38] in this head-fixed preparation to record large neural populations in PFC and other brain regions that have been implicated in ARD. Interestingly, the elevated quinine consumption that we observed in head-fixed mice may prove helpful in these electrophysiology studies by possibly strengthening neural encoding of the aversive stimulus. In addition to quinine adulteration, we will pursue alternative punishments that can be probabilistic and delayed in relation to drinking, such as tail shock, air puff, or optogenetic stimulation of peripheral nociceptive cells [80]. We believe these alternative aversive stimuli will allow for assessment of how deficits in neural encoding of delayed punishments may contribute to ARD. Overall, these future studies will provide simultaneous data on the precise neural firing patterns and behaviors associated with ARD across multiple brain regions, which will allow for advanced analyses of the neurocomputational mechanisms underlying ARD [81].

## Declarations of interest

none

## Notes

Support: This work was supported in part by NIH grant numbers: AA028265 (N.M.T.), AA007611 (C.C.L.), and AA029409 (C.C.L.).

https://figshare.com/articles/dataset/Data_and_Analysis_Code_Non-consummatory_behavior_signals_predict_aversion-resistance_in_head-fixed_mice/23546517

### Competing Interest Statement

The authors have declared no competing interest.

### Summary of Updates

Address comments from reviewers during peer review.

